# Autophagy-related genes *atg7* and *beclin1* are essential for energy metabolism and survival during the larval-to-juvenile transition stage of zebrafish

**DOI:** 10.1101/666883

**Authors:** Suzan Attia Mawed, Jin Zhang, Fan Ren, Jie Mei

**Author notes:** These authors contributed equally to this work (Suzan Attia Mawed, Jin Zhang).

## Abstract

High mortality is usually observed during the transition from larvae to juvenile in teleost which is related to the transition from endogenous to exogenous feeding. Autophagy is an evolutionary regulated cellular mechanism highly conserved in eukaryotic organisms to maintain energy homeostasis against stress including starvation. To investigate whether autophagy plays a role during the larval-juvenile transition, we generated *atg7* and *beclin1* zebrafish mutant lines using CRISPR/Cas9 technology. In this study, both *atg7* and *beclin1* null zebrafish exhibited a normal body confirmation; nevertheless, they completely died around 15 dpf and 9 dpf respectively. During larval-juvenile transition period, *atg7* and *beclin1* mutants were unable to cope with the metabolic stress after yolk absorption at 5 dpf and fail to activate autophagy in response to nutrient restriction, and without external feeding, all mutants died nearly at 8 dpf. Dramatic defects in the intestine architecture and metabolic functions in the liver were observed even though providing larvae with an external food supply, suggesting that autophagy isn’t only important during yolk depletion but also within food plenty. Treatment with rapamycin, an activator of autophagy, could effectively extend the survival time of both *atg7* and *beclin1* null zebrafish through lowering the metabolic rate while it couldn’t activate autophagy in mutants via the canonical pathway. Our findings provided a molecular evidence for the physiological, histological and metabolic changes that occur during the transition process from the larval to the juvenile stages and the chief role of autophagy on the body metabolism during these turning milestones.

**Author summary:** Zebrafish ***Danio rerio*** has emergrd one of the most powerful research models for studying genes expression during early embryogenesis and postnatal development. On the basis of the cell mechanisms, Macroautphagy, a natural regulated pathway disassembles unnecessary or dysfunctional components orchestrated by more than 36 autophagy related-genes conserved from yeast to mammals. Among those genes are *atg7* and *beclin1* which have been proved to play an important role in regulating post natal development in some mammals however their roles during zebrafish development still unedited. During this research, CRISPER/CAS9 were adopted to know *atg7* and *beclin1* knockout effects on the mutants’ metabolism during shifting from maternal yolk acquisition to exogenous feeding and the role of autophagy during the larvae to pre-juvenile development. Herein, we found out that larvae couldn’t abandon autophagy in both fasting and feeding conditions as larvae died earlier before pre-juvenile development despite feeding declaring the importance of autophagy not only to provide the cell with essential nutrients during starvation but also to get rid of cargos inside the eukaryotic cells. Briefly, if the larvae didn’t recycle those cargos due to autophagy perturbations, they will die despite providing suitable conditions including food and acclimatization.

## Introduction

On the basis of the developmental events, the transition from the larval to the juvenile stage is crucial during fish ontogeny since embryos shifting from maternal yolk acquisition to extrinsic food resources depending on the cellular metabolism and energy (1). Larva to juvenile transition is often rapid and involves many morphological, physiological, molecular, and behavioral adaptations which are energetically costly to suit the new environment (2). However, the molecular signals controlling the larval-to-juvenile transition remain largely unclear (1, 3).

Macroautophagy here referred to as autophagy, alters both anabolic and catabolic processes with the help of two key sensors, the activated protein kinase (AMPK), a key energy sensor and the mechanistic target of rapamycin (TOR), a key nutrients sensor(4). Moreover, autophagy involves in the cytosolic rearrangements needed for differentiation and growth during embryonic development, which can mediate protein and organelles turnover within a few hours (4, 5). Under the nutrient-rich condition, mTOR is activated and stimulates anabolic processes such as gluconeogenesis, protein synthesis, and energy metabolism, whereas catabolic pathway via autophagy is prohibited (6). During starvation stressors mTOR get switched off, thereby enabling the activation of autophagy in wild type zebrafish larvae after maternal yolk depletion at 6 dpf (7, 8). Moreover, reactivation of mTOR attenuates autophagy and initiates lysosome regeneration (9).

Up to date, several studies reported that *atg7* silencing results in neurodegeneration (10), histopathological changes in the liver of mice (11, 12), and tumorigenesis in both murine and human (13). In addition, *atg7* accelerates hepatic glucose production through the action of glucogenic amino acids in the hepatocytes of the null mouse during nutrient depletion (14).On the other side, homozygous deletion of *beclin1* is lethal in mice embryo while *beclin1*^+/−^ embryos suffer from a high incidence of spontaneous tumors (15). The novel role of the autophagy pathway in intestinal microbiota and innate immunity in the small intestine was revealed using mice with mutations in *atg16L1*, *atg5*, and *atg7*(16, 17). Autophagy is necessary to maintain the unique structure and proper functions of the exocrine pancreas and trypsinogen activation (18, 19).

In this study, we used *atg7* and *beclin1*mutant zebrafish as models to investigate the relationship between autophagy and metabolism during zebrafish development. There were no phenotypes in *atg7*^−/−^ and *beclin1*^−/−^mutant zebrafish during embryonic development however both *atg7* and *beclin1* mutant strains have darken prominent liver in vivo during the period of larval-to-juvenile transition and died within 15dpf and 9dpf respectively. A*tg7* and *beclin1*mutants were unable to cope with the metabolic stress after yolk absorption at 5dpf and fail to activate autophagy in response to nutrient restriction resulted in perturbations in hepatic glycogen/lipid metabolism. Collectively, our results suggest that autophagy-related genes *atg7* and *beclin1* are required for maintaining energy homeostasis and mediates liver metabolism during this transition state.

## Results

### Generation of atg7 and beclin1 mutagenesis by CRISPR/Cas9

CRISPR/Cas9 system was conducted to generate our zebrafish mutants. Briefly, the sgRNA targeting sites were chosen in the 10^th^ and 4^th^ exons of *atg7* and *beclin1*, respectively. Finally, two *atg7* mutant lines were established, which contained 5-bp and 2-bp deletion, respectively, named *atg7*Δ5 and *atg7*Δ2 (Fig.1A). Zebrafish *atg7* protein consists of two functional domains including ATG-N superfamily and E1 enzyme superfamily. The deletions in *atg7*Δ5 and *atg7*Δ2 resulted in a frameshift that caused premature stop codons (Fig.1B). On the other side of genetic mutation, one *beclin1* mutant line was established, which contained 8-bp deletion, named *beclin1*Δ8 (Fig. 1C). Zebrafish *beclin1* protein consists of two functional domains including BH3 and APG6 superfamily. The deletions in *beclin1*Δ8 resulted in a frameshift that caused premature stop codons (Fig.1D).

**Figure1.**
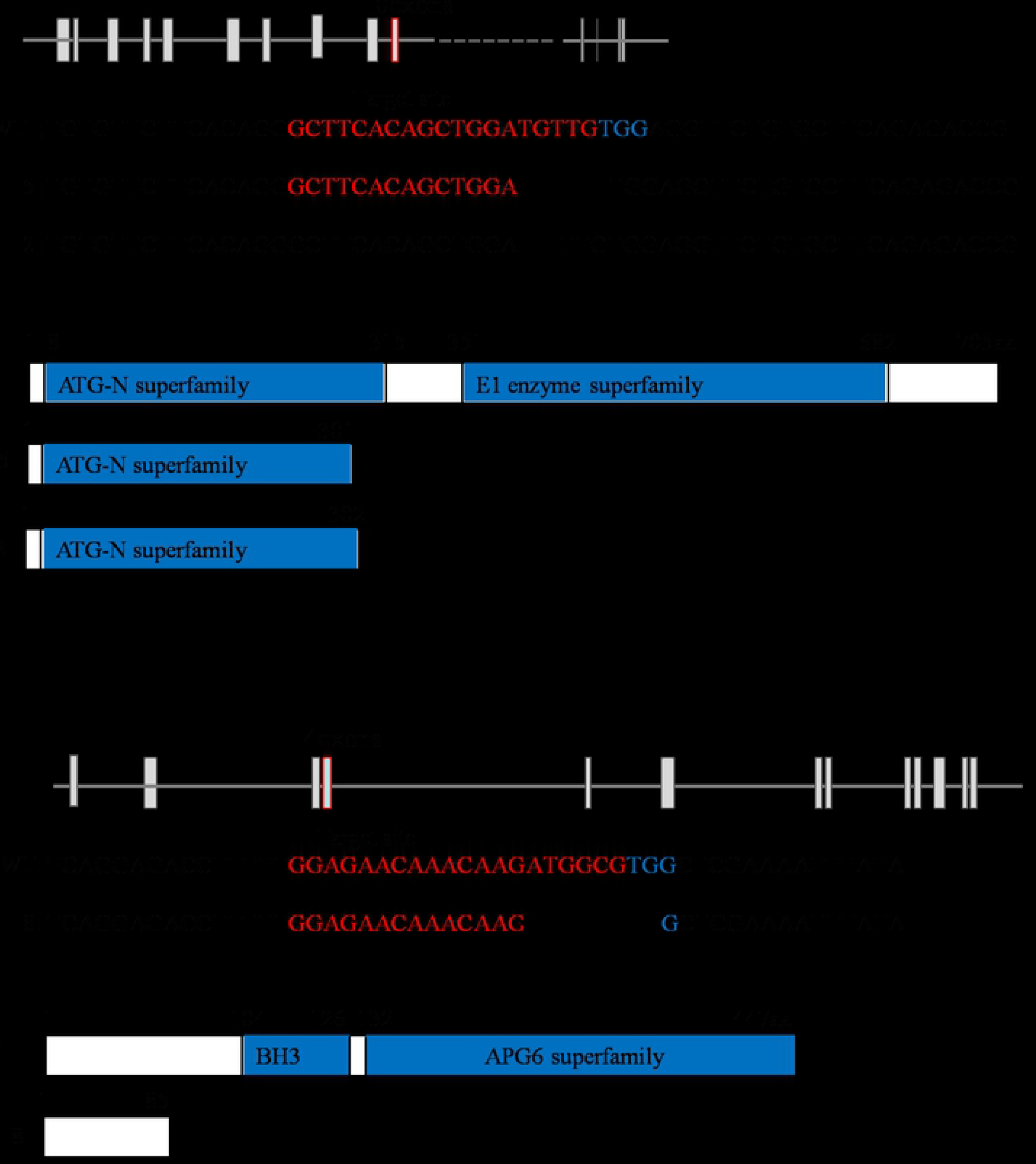
Generation of *atg7* and *beclin1* mutated zebrafish: (A): Schematic representation of zebrafish *atg7* and its two mutated types (*atg7Δ5* and *atg7Δ2*). The thin lines and grey boxes represent the introns and exons, respectively. The sgRNA target sequence is shown in red, followed by a PAM sequence “TGG” shown in blue. (B): Illustration of deduced protein structures of wild-type *atg7* (top) and two mutageneses (middle and bottom).(C): Schematic representation of zebrafish *beclin 1* and mutant line (*beclin1Δ8*). The sgRNA target sequence is shown in red, followed by a PAM sequence “TGG”.(D): The structure of the deduced protein domain of wild type *beclin1* (top) and the represented mutant line (bottom).

### Loss of zebrafish atg7 or beclin1 function results in lethality during the larval-to-juvenile transition

When crossing the male and female *atg7*^+/−^ or *beclin1*^+/−^ zebrafish, no obvious defects were observed during the early embryonic development. Interestingly, *atg7* and *beclin1* mutants exhibited darken prominent liver compared with the wild type and heterozygous strains of the same population during larvae development (Fig.2A). To clarify this issue, whole mount Oil red O (ORO) staining was carried out at 7 dpf that ensured the lipid accumulation in the *atg7* mutant liver and in the whole gastrointestinal tract (GIT) of *beclin1* littermates, while wild type and heterozygous larvae have nearly the same phenotype of null lipid accumulation (Fig.2B). In zebrafish, the maternal yolk almost depleted after 5-6 dpf without exogenous feeding WT zebrafish larvae began to die as early as 6-7 dpf and completely died as later as 13 dpf. Meanwhile, *atg7* and *beclin1* homozygous mutants died from 6 to 9 dpf and 6 to 8 dpf respectively, which were identified by the phenotype and genotyping (Figs. 2C, 2D). Moreover, exogenous feeding couldn’t rescue the life of *atg7*^−/−^ or *beclin1*^−/−^ zebrafish as they started to die from 6 to 15 dpf and 6 to 9 dpf, respectively (Figs. 2E, 2F). Since *atg7Δ5* and *atg7Δ2* have nearly the same knockout effect on the protein domain, and survival rate, *atg7Δ5* was applied as an example of *atg7* knocking out in our further study.

**Figure2.**
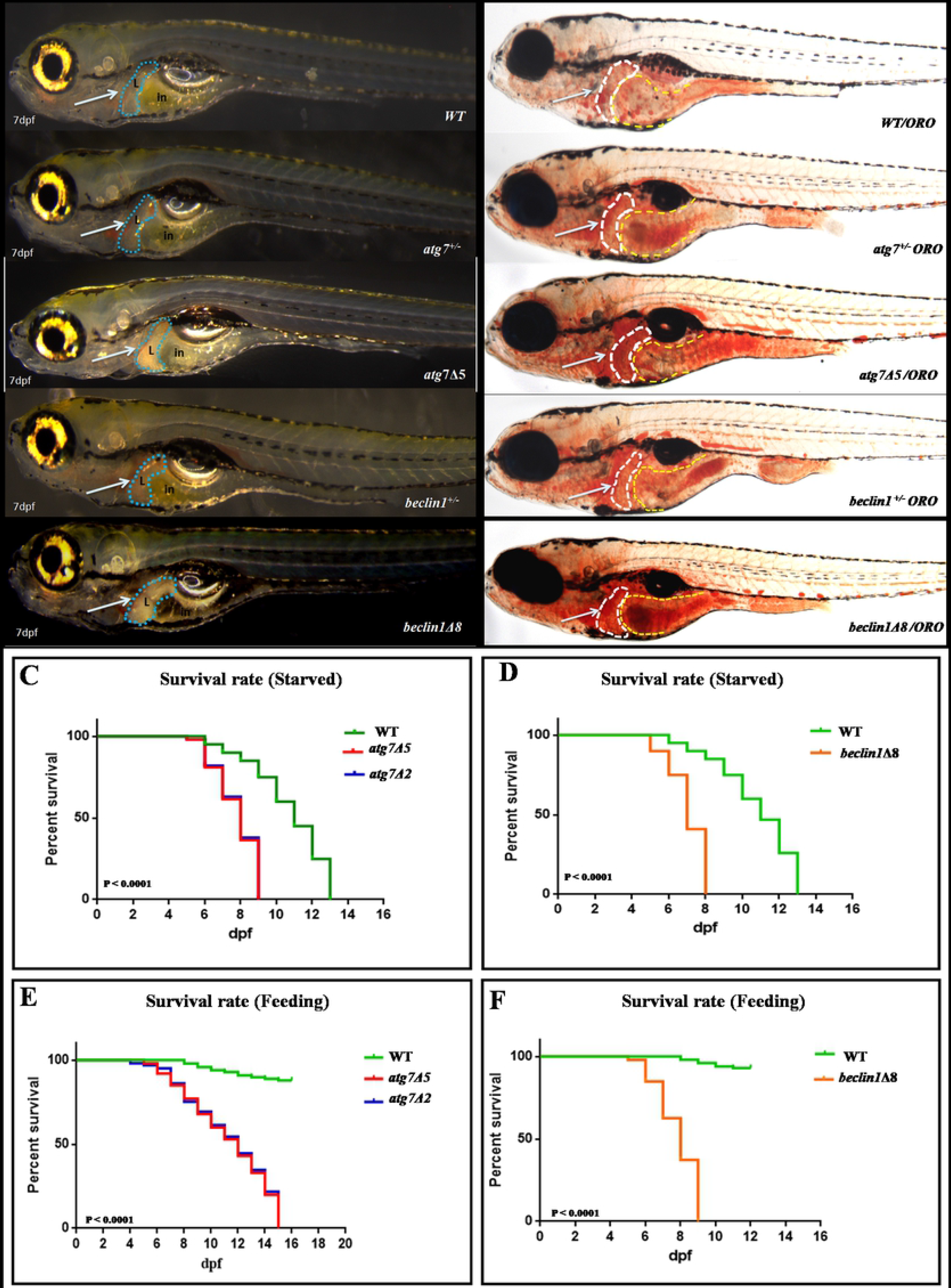
Mutations of *atg7* and *beclin1* caused lethality during the larval-to-juvenile transition. (A): 7dpf larvae of the indicated genotypes shown live.(B): Whole-mount Oil Red O staining indicates lipid droplets stained in-situ of 7dpf genotyped larvae. Arrows indicate zebrafish liver with no detected defrence between wild type and heterozygous of both strains. (C-D): Meire Kaplan graphs depicting the surviving rate upon fasting,WT started to die at 6 dpf along with the two mutants lines of *atg7*^−/−^(Fig.C) and *beclin1Δ8* (Fig.D). WT could survive till 13 dpf whereas *atg7*^−/−^ and *beclin1*^−/−^ completely died at 9 and 8 dpf respectively.(E-F): Meire Kaplan graphs depicting the surviving rate upon feeding of two mutant lines of *atg7* (*atg7Δ5* and *atg7Δ2*) and *beclin1Δ8* compared with WT.

During starvation in wild-type zebrafish larvae, mRNA expression of autophagy-related genes including *atg7*, *beclin1*, *atg5*, *atg12* was gradually increased during starvation from 5 dpf to 8 dpf and began to reduce upon feeding, whereas *p62* expression was gradually reduced upon starvation and recovered after feeding, indicating that autophagy was induced during the yolk transition state (Figs. 3A-3E). To further validate the loss of *atg7* and *beclin1* function in mutant zebrafish; qRT-PCR was conducted at 5^th^ and 7^th^ dpf on starved mutant larvae and at 8^th^ dpf on the fed group to detect the mRNA expression of the previously mentioned autophagy-related genes in mutants compared with their wild type siblings (Figs. 3F-3J). Interestingly, *atg7* and *beclin1* null larvae exhibited a very low gene expression of *atg7* and *beclin1* respectively (Figs.3F, 3G). Moreover, *atg5* and *atg12* genes expression showed no big difference during the transition period (Figs. 3H, 3I). On the other side, in both mutants, *p62* expression within the transition period exhibited a significant increase despite feeding at 8 dpf (Fig. 3J). Our results suggest that *atg7* and *beclin1* are essential for the larval-juvenile transition around 5-14 dpf and indicates the up-regulation of the autophagy process upon yolk termination that was prohibited in the mutants of our study.

**Figure3.**
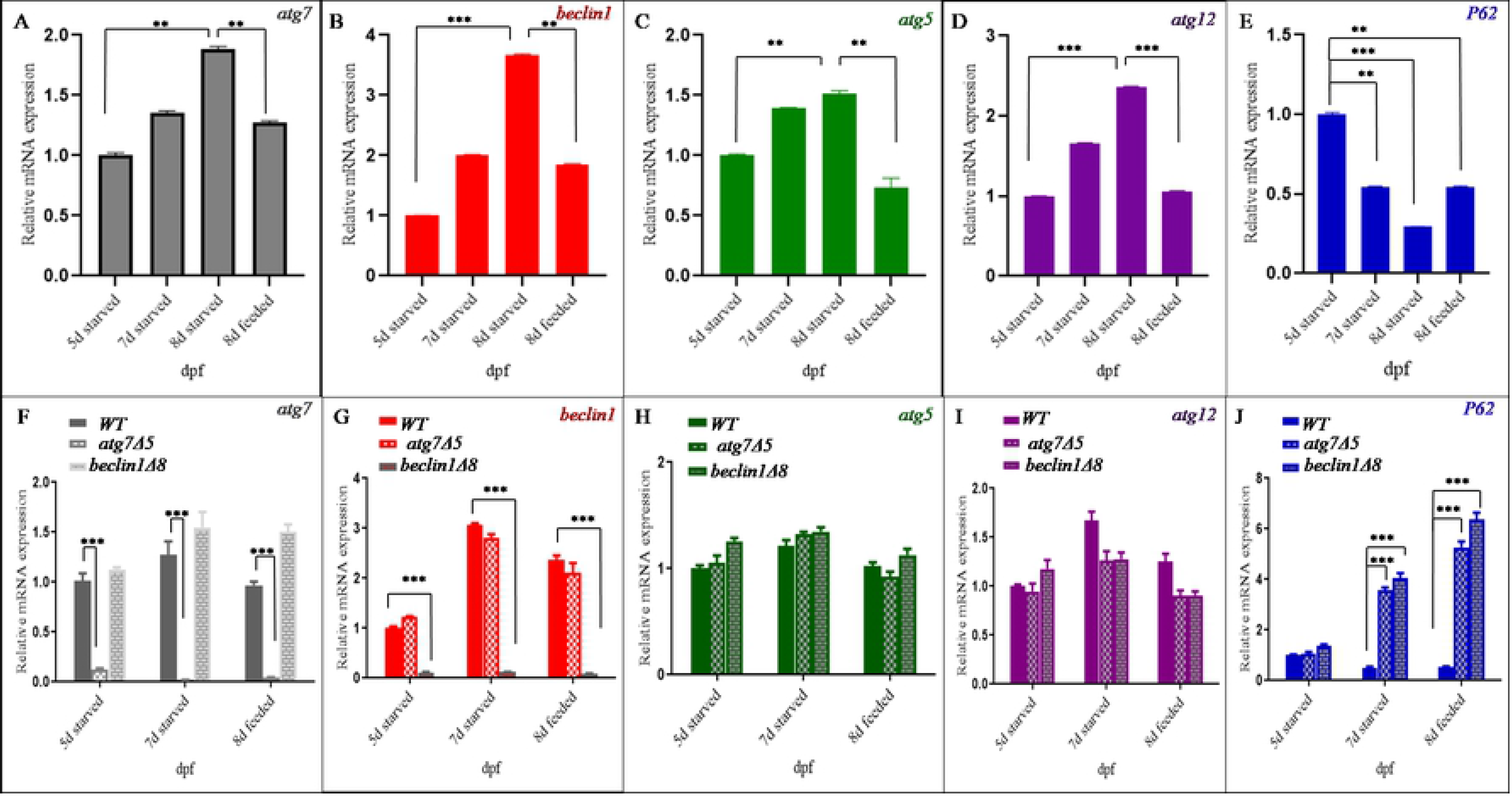
Wild type but not mutants exhibited autophagy induction during larval-juvenile transitionm: (A-E) mRNA expression of *atg7*, *beclin1*, *atg5*, *atg12*, *and p62* repectively were evaluated by qRT-PCR in wild-type indicated actual autophagy induction upon starvation (5-8dpf). (F-J): Characterization of zebrafish *atg7* and *beclin1* mutations through mRNA expression using previous autophagy-related genes compared with their wild type siblings at 5,7 and 8 dpf with a significant lower expression of knockedout genes and *p62* elevating. Data were representative of three independent experiments and expressed as mean ± SD.* P <0.05, * *P<0.01, and * * *P < 0.001.

### Gastrointestinal tract (GIT) developmental defects displayed during the weaning period between the endogenous and exogenous feeding in atg7- and beclin1-null embryos

Since the transition from the larval to the juvenile stage is also a transition from endogenous to exogenous feeding, whole mount in situ hybridization was conducted to determine whether the early death of *atg7* and *beclin1* mutants was due to developmental defects or metabolic inadequacy. *Fabp10a*, a marker of the liver (Figs. 4A-4C) and *Villin1* a marker for intestine (Figs. 4G-4I) demonstrated a normal expression pattern before maternal yolk depletion at 72hpf and 5dpf, respectively between WT and *atg7* or *beclin1* mutants, at that time embryos still depend on the maternal nutrition. Upon yolk depletion at 6 dpf, there was a remarkable decrease in the gene expression in the mutant strains compared with their WT counterparts for *fabp10a* (Figs. 4D-4F) as well as *Villin1* expression was down-regulated (Figs.4J-4L), suggesting the beginning of the malformation during the transition state which might be responsible for the early death of mutant strains.

**Figure4.**
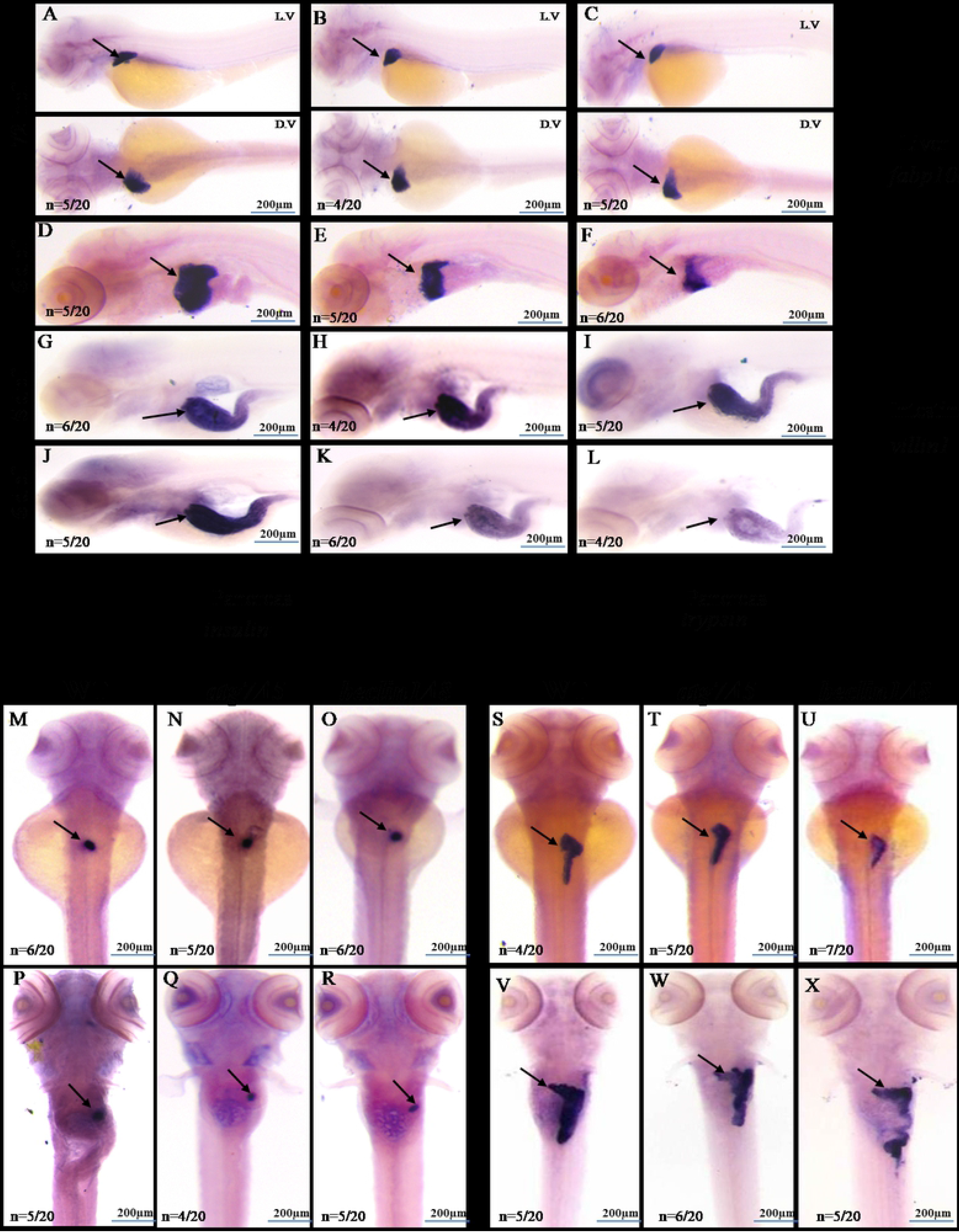
*Atg7* and *beclin1* affect the gastrointestinal tract and accessory organs after yolk depletion. RNA in situ hybridization was performed with *fabp10a* (A-F), *villin*. (G-L), showing lower signals for liver and enterocyte differentiation respectively at 6 dpf. (M-R) pancreatic *insulin*, a marker of the endocrine pancreas at 72 hpf and 6 dpf. (S-X): larvae stained with probes against *trypsin*, a marker for terminal differentiation of the exocrine pancreas that exibited at lower expressionin mutrant *atg7* and *beclin1*. Number of larvae shown within the whole poulation number (n=20) L.V: laterl view, D.V: dorsal view.

To further clarify the role of autophagy in pancreatic activity, we examined *trypsin* expression for exocrine activity and *insulin* expression for the endocrine pancreas. Loss of *atg7* and *beclin1* function didn’t directly interfere with the *insulin* expression at the early development as it exhibited differences neither prior nor after yolk depletion from 72 hpf till 6 dpf (Figs.4M-4R). On the other side, *trypsin*, a digestive enzyme gene expressed in the exocrine pancreas, displayed a lower signal in mutant *atg7* and *beclin1* comparing with WT at 72 hpf (Figs.4S-4U) and with more remarkable down-regulation at 6 dpf (Figs.4V-4X). In wild-type zebrafish, trypsin synthesized normally and drained into the duodenum during the weaning period. Contrarily, it was expressed in a reduced signal area in the *atg7* as well as *beclin1* mutants. After photos capturing, all larvae were genotyped to confirm our results which indicated that autophagy dysfunction symptoms manifested in the later developmental stages during the weaning period between endogenous and exogenous feeding suggesting a critical window during the larval-juvenile transition.

### Atg7 and beclin1 mutants fail to reactivate autophagy in response to starvation

In wild type during the transition period (5-7 dpf), the maternal yolk was depleted and mTOR was switched off enabling the activation of autophagy (20). Since autophagy activity is commonly detected by higher accumulation of autophagy microtubule-associated protein 1 light chain 3, LC3-II (21) and degradation of ubiquitin-conjugating protein P62/SQSTM1 (22), the two markers were used to monitor the autophagy flux. In contrast to higher LC3-II protein accumulation in the liver and intestine of WT larvae (Fig. 5A a), both *atg7* and *beclin1* mutants showed a great reduction in LC3-II expressions (Figs. 5A b, c). On the other side, P62/SQSTM1 expression was elevated in mutants especially *beclin1*compared with their WT siblings (Figs. 5A d-f).The same results were obtained from the western-blot analysis of LC3A/B that usually exhibited in two bands: LC3-I and LC3-II. LC3-II is more appropriate for detecting autophagy since it ensures the complete transformation of pro-LC3-I. Herein, we detected a dysfunction in the autophagic signaling cascade in the mutant strains at 7 dpf via accumulation of P62/SQSTM1 and reduction of LC3-II versus their wild type littermates (Fig. 5B). For further verification, immunofluorescence of liver sections from WT and both mutants also confirmed the latter results (Figs.5C, 5D). Accordingly, our results further suggest that *atg7* and *beclin1* mutants fail to activate autophagy under nutrient depletion as well as the predicted correlation between autophagy disruption and GIT metabolic disturbance.

**Figure5.**
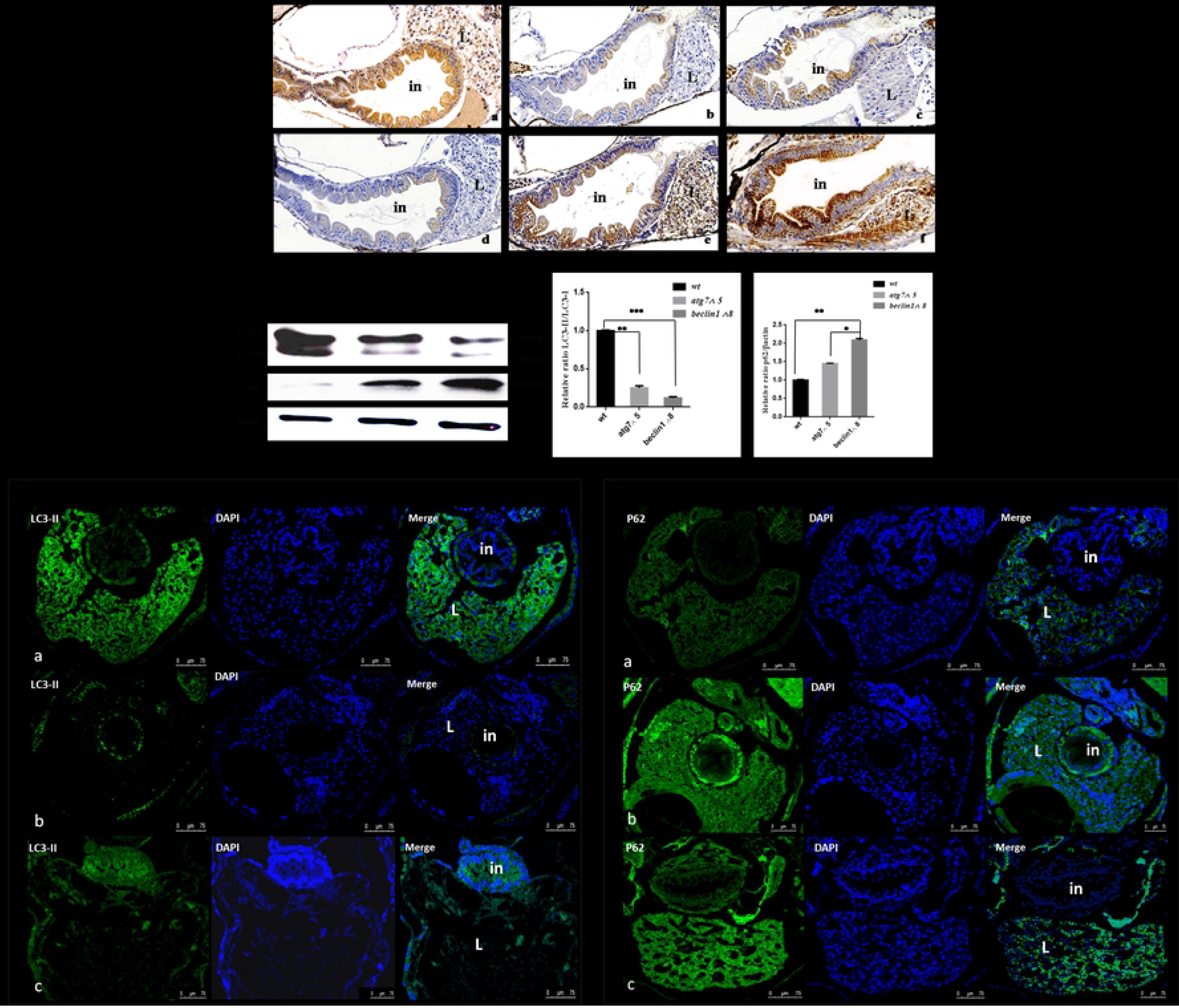
*Mutant strains were* unable to activate autophagy during metabolic stress. (A): Immunohistochemistry assay of longitudinal sections (5 μm) of starved 7 dpf larvae blocked against LC3-II (a-c) and P62/SQSTM1 (d-f) showed impairment autophagy flux in both mutants unlike wild type. (B): Representative western blots of the three genotyped strains showed relative protein expression of P62/SQSTM1 and LC3-II, β-actin was used as the control. The ratio of p62/β-actin and LC3-II/β-actin quantified by NIH software Image J. Progressive P62/SQSTM1 accumulation in mutant larvae reveals the impaired autophagy and liver toxicity. (C-D): Immunofluorescence of the hepatic transverse regions of both *atg7* and *beclin1* mutants and wild type confirming the autophagy blocking in the mutants. L: liver; in: intestine. *P < 0.05, **P < 0.01, and ***P < 0.001.

### Loss of zebrafish atg7 or beclin1 function leads to abnormal intestinal architecture after maternal yolk consumption

In order to investigate the relationship between autophagy activity and intestinal architecture, longitudinal sections of 7 dpf and 14 dpf larvae were carried out. At 7 dpf, intestinal bulb of WT larvae was completely developed with normal folds in the intestinal epithelium and visible villi (Fig.6A). On the other side, mutant intestinal epithelium seems to be disorganized (Figs.6B, 6C).The lining epithelia of *atg7*^−/−^ and *beclin1*^−/−^ appeared abnormal with pyknotic or loss of nuclear polarity and cellular structure that led to disturbance of the intestinal barrier and absorptive functions. By 14 dpf, WT intestinal mucosa consists of columnar enterocytes with basal arranged nuclei (Fig. 6D). On the contrary, *atg7* mutants exhibited abnormal villous architecture, microvillus atrophy and impaired proliferation of intestinal epithelial cells accompanied by hyperplasia that often leads to extensive folding and formation of pseudo-crypts (Fig.6E). As expected, *beclin1*^−/−^ zebrafish exhibited rapid dramatic intestinal changes than in *atg7*^−/−^ zebrafish, suggesting that *beclin1*^−/−^ zebrafish at 7 dpf were in advanced idleness made them comparable to *atg7*^−/−^ zebrafish at 14 dpf. Briefly, *beclin1* mutants showed more severe loss of intestinal epithelium structure and function with defective villi formation that appeared earlier than *atg7*^−/−^ zebrafish.

**Figure6.**
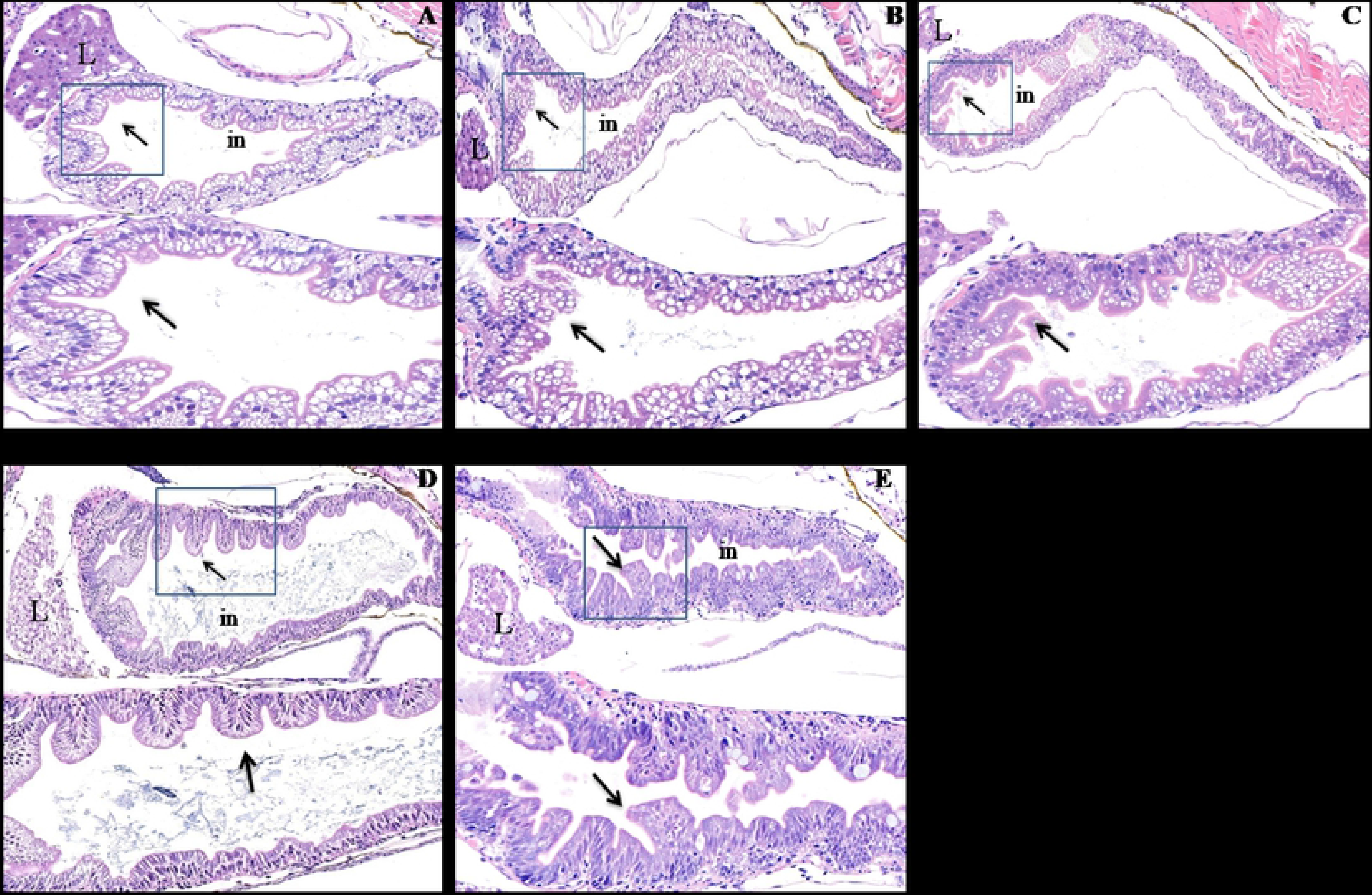
Loss of *atg7* and *beclin1* led to defects in the gastrointestinal architecture: (A-E): Representative photomicrographs of H&E stained sections of 7dpf and 14 dpf larvae demonstrating the disproportionate intestinal architecture of *atg7Δ5* and *beclin1Δ8* compared with WT which exhibited nicely organized villi with linearized enterocytes and arranged nuclei (A and D) unlike the improper pseudo villi and nuclear atypia in mutants (B, C and E).The lower pictures are magnified part of the intestinal villi inside the blue squares.

### Zebrafish atg7 or beclin1 knockout associates with liver metabolic disequilibrium

During the process of endogenous-exogenous transition, liver plays a vital role in the regulation of systemic glucose and lipid fluxes during feeding and fasting. After yolk depletion, liver provides glucose for all tissues of the body by breaking down its own stores of glycogen via the process of glycolysis (23, 24). At the same time, liver replaces the consumed glycogen through gluconeogenesis and/or lipolysis that includes the formation of alternative glycogen from non carbohydrate sources, such as lactate, pyruvate, glycerol, and alanine (25, 26).

Herein, liver glycogen content was detected by periodic acid-Schiff (PAS) staining before and after yolk absorption, at 5, 7 and 14 dpf for *atg7*^−/−^and at 5, 7 dpf for *beclin1*^−/−^mutants. At 5 dpf, wild-type and both mutant strains contain an adequate amount of hepatic glycogen (Figs.7A, a-c). At 7dpf, unlike WT, the hepatic glycogen was gradually depleted during the transition state in mutant strains which suggested an increase of glycolysis and decrease of gluconeogenesis (Figs.7Ad-f). Within 14 dpf, glycogen stock in *atg7*-null hepatocytes totally vanished compared with WT (Fig.7A g, h). Interestingly, *atg7*^−/−^ intestine still contained residual glycogen suggesting indigestion as liver glycogen shed towards the intestine. However, it neither digested nor absorbed due to a disturbance in the trypsin secretion and disordered intestinal epithelia.

**Figure7.**
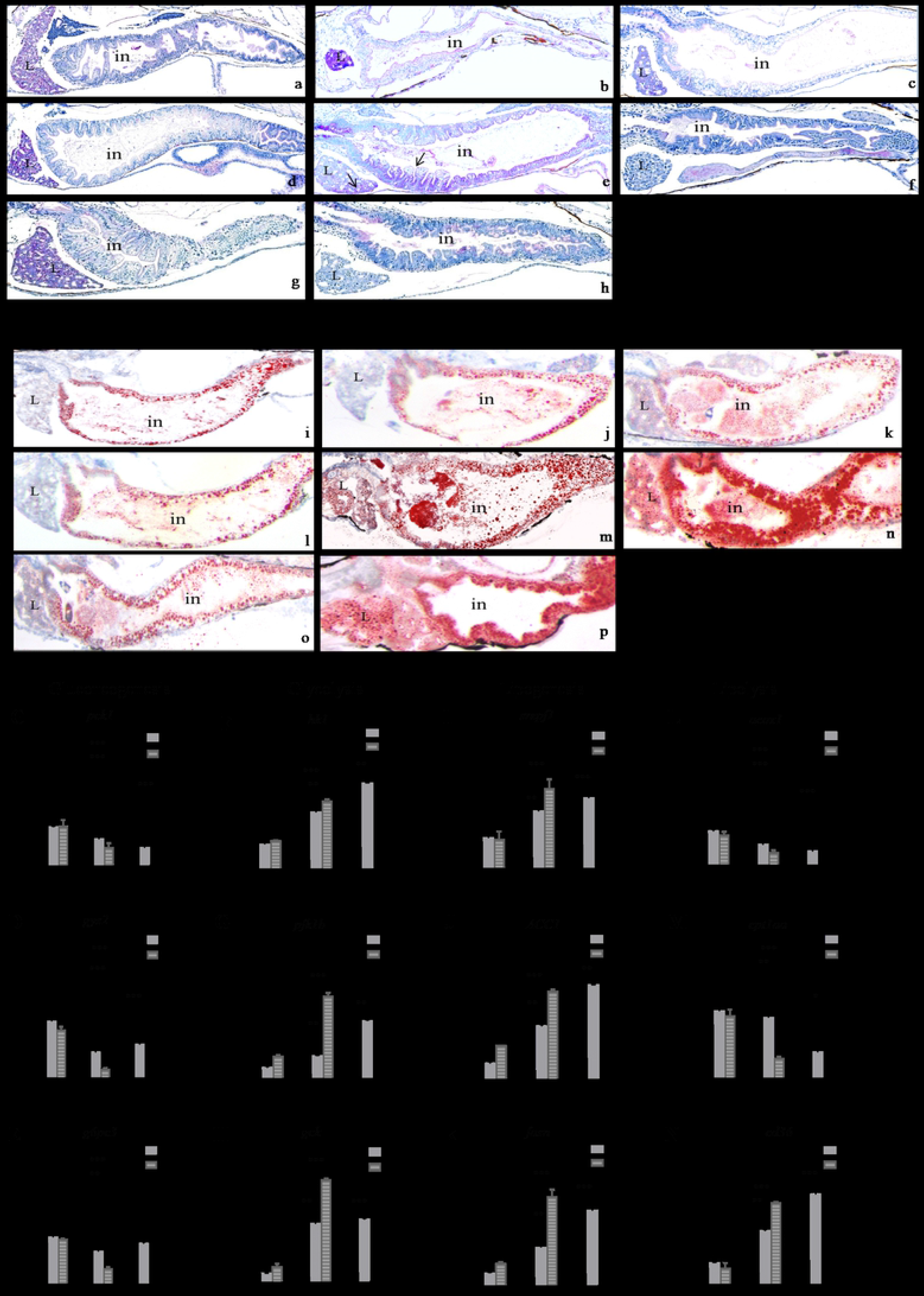
*Atg7* and *beclin1* mutant exhibit disturbed hepatic hallmarks and affect glycogen-lipid flux in response to starvation. (A): Longitudinal sections stained with PAS of WT, *atg7* and *beclin1* mutant livers at 5, 7 and 14 dpf. WT liver still contains an adequate amount of glycogen from the 5 dpf till 14 dpf (a, d, g), whereas glycogen depleted gradually in *atg7Δ5* and *beclin1Δ8*. At 7dpf, *atg7Δ5* showed disturbance of hepatic glycogen absorption represented by black arrows (7A e). *Beclin1* mutants exhausted earlier where hepatic glycogen vanished by 7dpf suggesting earlier metabolic disturbance (7A f). (B): Oil Red O staining of longitudinal frozen sections passed through the liver with remarkable undigested intestinal lipid and triglycerides aggregation at 5 and 7dpf in mutant livers, suggesting steatohepatitis that appeared earlier in *beclin1* mutants (7B n) than *atg7* mutants (7B m). (C-E): mRNA expression of genes involved in hepatic gluconeogenesis. (F-H): mRNA expression of selected genes involved in glycolysis process depicting the relation between autophagy knockout and glycogen depletion during feeding-fasting transition. (I-K): mRNA expression of genes involved in lipogenesis. (L-N): mRNA expression of genes involved in lipolysis indicating hepatic steatosis in *atg7* and *beclin1* null larvae via inhibition of fatty acids β oxidation. Data were representative of three independent experiments and expressed as mean ± SD.* P <0.05, * *P<0.01, and * * *P < 0.001. L: liver. in: intestine.

Lipids can be visualized with the Oil Red O (ORO) dye. Compared with 5,7 and 14 dpf WT larvae (Figs. 7 B i, l, o), *atg7*-and *beclin1*-null larvae displayed abnormal retention of lipids in the liver, as well as undigested fat droplets inside the intestine accumulated due to steatosis (Figs. 7B m, n, p). *Beclin1* mutants exhibited accelerated morphological and cellular hallmarks of starvation at 7 dpf since hepatic lipid couldn’t be utilized in regard to autophagy deficiency in the mutant liver (Fig.7B n).

To further investigate the relationship between autophagy activity and glycolipid flux, mRNA expression of genes involved in hepatic glycogen and lipid metabolism were evaluated before and after yolk transition at 5, 7 dpf during starvation and at 14 dpf in feeding condition. Starvation till 7 days will give an indication about the liver metabolic equilibrium during fasting before shifting to exogenous feeding after 7 dpf till 14 dpf. In our study, genes involved in gluconeogenesis including *pck1*, *gys2*, *and g6pc3* (Figs.7C-7E) showed no significant difference between the three studied groups at 5dpf, however, they increased upon starvation at 7 dpf and decreased again after feeding at 14 dpf in wild type. Interestingly, those genes couldn’t be revived in both *atg7* and *beclin1* mutants during the period of starvation at 7 dpf compared with their wild type siblings. Contrarily, the mRNA expression of *hk1*, *pfk1b*, *gck* genes involved in glycolysis (Figs.7F-7H) were remarkably elevated in both mutants compared with their WT within the same population indicating a disturbance in glycogen synthesis and induction of glycolysis or glycogen consuming during the starvation in both mutants without renewing even though feeding *atg7* mutants till 14 dpf.

In another point of view, genes involved in lipid metabolism were also quantified in the same timeline of glycogen metabolism, herein, genes involved in lipogenesis process including sterol regulatory element binding transcription factor 1*(srepf1)* and other lipogenic enzymes such as acetyl-CoA carboxylase 1 *(ACC1)* and fatty acid synthase *(fasn)* were highly induced in mutants versus wild type (Figs.7I-7K). Interestingly, these genes still highly expressed in mutants of *atg7* at 14 dpf despite feeding. On the other side, the expression of genes responsible for hepatic fatty acid β-oxidation including *acox1*, *cpt1aa* (Figs.7L, 7M) declined sharply in the same mutants confirming hepatic steatosis via lipid accumulation. For further description, Fatty acid translocase *(FAT/cd36)*, a membrane protein participated in fatty acid uptake of hepatocytes, was significantly increased in both mutant strains at 7 dpf and at 14 dpf of *atg7* mutants (Fig.7N). Briefly, autophagy perturbation was highly correlated with the induction of glycolysis and lipogenesis and inhibition of gluconeogenesis and lipolysis during the larval-juvenile transition of null *atg7* and *beclin1*.

### Rapamycin enhanced the survival rate of mutants via yolk retention but not through autophagy induction

Pharmacological treatment via rapamycin induced a mild general developmental delay and could slow down the dramatic deterioration in mutant strains resulted in increasing the lifespan 2-3 days in both *atg7* and *beclin1* mutants (Figs.8A, 8B). At 7 dpf, we tried to focus on the effect of rapamycin on the metabolism as well as autophagy during this time point. Accordingly, mutants of control group exhibit dark liver compared with the wild type, while treated groups manifested significant delay in yolk absorption (Fig. 8C). Histological observations confirmed the growth retardation along GIT as intestine of treated larvae were barely detected and yolk sac occupied almost all the abdominal cavity. Interestingly, we found out that rapamycin could enhance the metabolism of both mutant strains by glycogen accumulation and hepatic lipid drainage similar to their WT littermates (Fig. 8D).

**Figure8.**
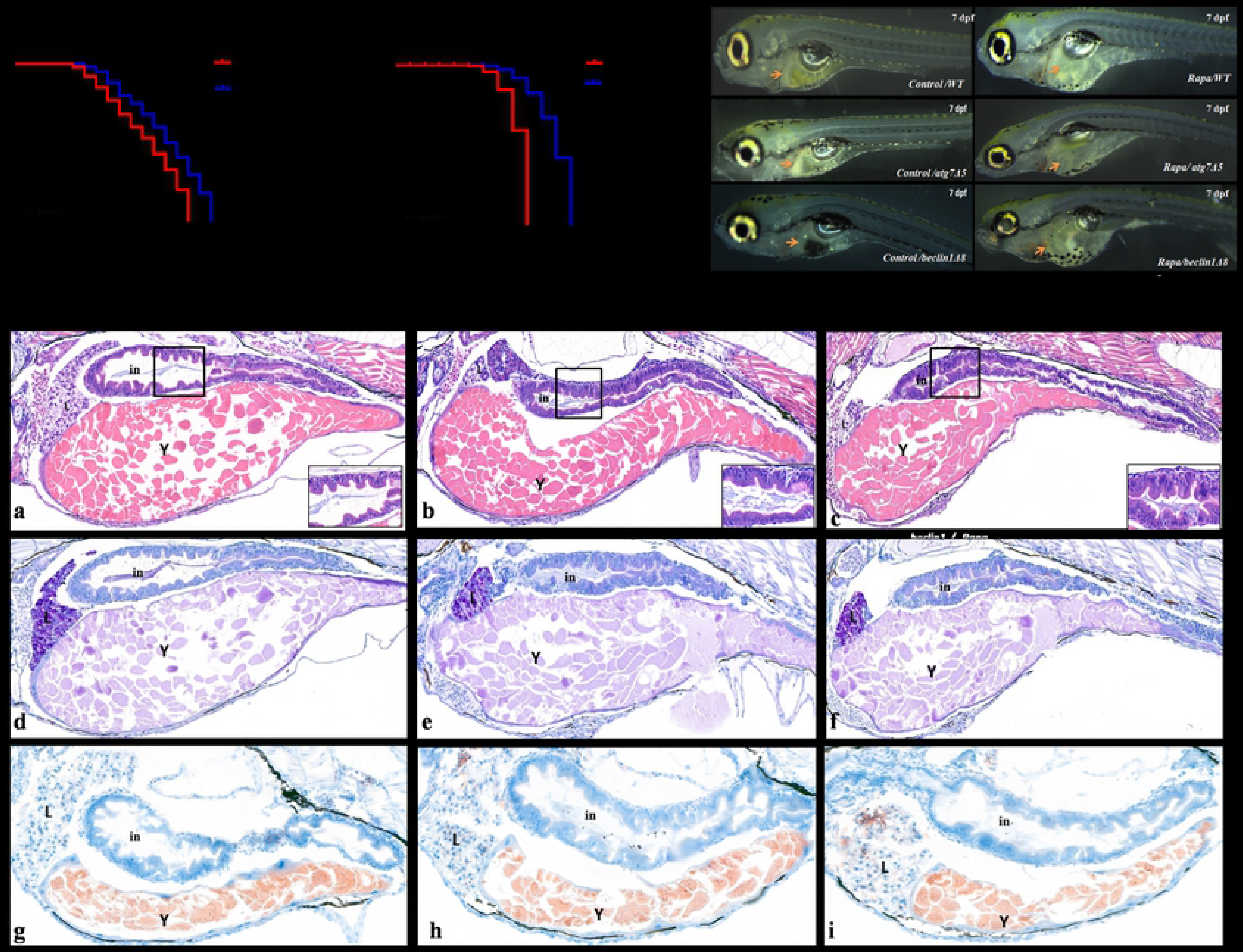
Rapamycin affects the survival rate and induced morphological developmental delay. (A-B): Meire Kaplan graphs depicting the survival rate after rapamycin treatment in *atg7* and *beclin1* mutants. (C): Control and treated embryos are shown live. Embryos treated with rapamycin have a generalized developmental delay with yolk retention and deceleration of digestive system growth. (D): Longitudinal sections of rapamycin-treated embryos stained with H&E (a-c), PAS (d-f) and ORO (g-i) indicating metabolic refreshment after rapamycin and intestinal development delay. in: intestine; L: liver; Y: yolk.

Treatment with rapamycin increased the protein levels of both LC3-II and P62/SQSTM1 in WT larvae (Fig. 9Ad, 9Bd).However, rapamycin had no obvious impact on autophagy induction in *atg7*- and *beclin1*-null larvae, since it couldn’t elevate LC3-II or restore P62/SQSTM1 (Fig. 9A e, f- 9B e, f). For further confirmation, protein expressions of LC3-I/II and P62/SQSTM1 were detected by western-blot analysis that ensures perturbation of Atg5/Atg7 dependent pathway and the disability of rapamycin for autophagy reactivation in both mutant strains (Fig. 9C). Collectively, rapamycin has no further effects on autophagy induction as it depends on mTOR and ATGs core that involved in the conventional autophagy pathway.

**Figure9.**
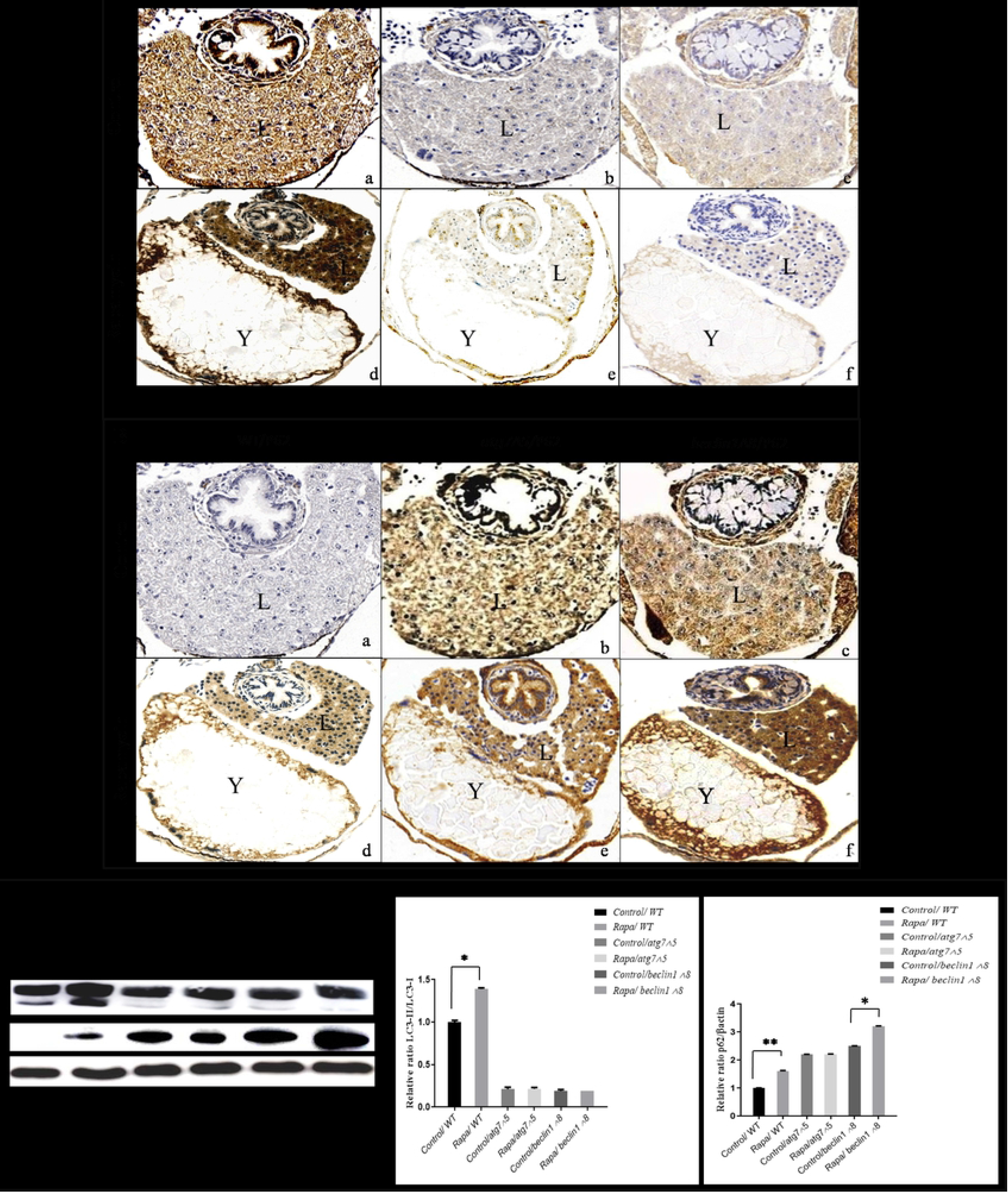
Effects of rapamycin on autophagy flux. (A): Immunohistochemistry assay of transverse sections from control and rapamycin treated larvae with no impact on LC3-II in mutants contrary WT. (B): Rapamycin couldn’t restore P62/SQSTM1 in both mutants as well as in WT. (C): Representative immunoblots showed LC3 I/II and P62/SQSTM1 proteins of control and rapamycin-treated embryos at 7 dpf starved. β-actin was used for normalization and relative protein levels were quantified using NIH software Image J. * P <0.05, * *P<0.01, and * * *P < 0.001. L: liver; Y: yolk.

## Discussion

Autophagy manipulated-deficient models give strong evidence on the role of autophagy in the early embryogenesis and postnatal development. Therefore, the metabolic balance between food restriction, autophagy and mTOR displays a dynamic pattern that affects the size of the organism, proper differentiation, and longevity (27, 28). As long as the food resource is available, the mammalian target of rapamycin (mTOR) allows cellular anabolism for building necessary blocks essential for growth and proliferation. During the larval-juvenile transition state when the maternal yolk depletes, larvae underwent starvation before shifting from endogenous to exogenous feeding, where they depend on catabolism and/or autophagy to compensate nutrient deprivation and maintain survival (29, 30). Surprisingly, unlike *atg7^−/−^* and *beclin1^−/−^*mice embryos that born healthy but died within one day neonatal (11, 15), *atg7* and *beclin1*, null zebrafish in our study exhibited a normal body confirmation, however, died around 14-15dpf and 8-9 dpf respectively suffering from reduced glycogen and hepatic steatosis. Furthermore, upon starvation, WT larvae could survive till 13dpf however, starved *atg7^−/−^* and *beclin1^−/−^*quickly finished yolk around 6 dpf and died within 7-8 dpf due to food restriction and blocking of the autophagy pathway. Similarly, loss-of-function mutations of *atg7* and *beclin1* reduce the life span of *Caenorhabditis elegans* (31). Our results authenticate the importance of autophagy during starvation as well as food richness.

Among all autophagy genes, *beclin1* and *atg7* have been focused in our study. Vesicle nucleation requires *beclin1* that forms a complex with the class-III phosphatidylinositol3-kinase (PI3K) Vps34. Moreover, the step of subsequent phagophore elongation requires *atg7* that controls the conjugation between the two ubiquitin-like conjugation pathways; *atg5-atg12* pathway and microtubule-associated protein1 light chain3 (LC3) lipidation to form membrane-associated LC3-II (32, 33).

Since liver, intestine, and exocrine pancreas are derived from the endoderm during the early embryonic development (34),WISH experiment reveals the down-regulation of gene markers related to those three digestive organs after 5 dpf. Generalized defects in the digestive derivatives indicate the importance of *atg7* and *beclin1* in the early endoderm differentiation. Interestingly, *trypsin* marker was substantially reduced that indicating its important role in the digestion process (35, 36). The down-regulation of *trypsin* secretion was also detected in *atg5*^−/−^ mice that were found to reduce pancreatic trypsin secretion and attenuate pancreatic damage (19).

For mutagenesis characterization, *atg7* and *beclin1* mRNA expressions were severely reduced in *atg7* and *beclin1* mutants, respectively indicating the effective gene knockout. Moreover, starvation till 7dpf elevated the expression of *p62* indicating autophagy disturbance in both mutants during the endogenous-exogenous transition, these findings put *p62* at a critical situation that control both cell death and survival(37). On the other side, the expression of *atg5* and *atg12* nearly didn’t change from their wild type siblings however *atg5–atg12* conjugation that is essential for autophagosome formation is impaired due to the absence of the *atg7* mediation in both mutants this also was previously explained elsewhere as *atg7* silencing resulted in loss of the *atg5–atg12* conjugate but doesn’t otherwise affect gene expression (38). On approach to evaluating autophagic flux, contrarily to wild type, our mutants show higher accumulation of p62/SQSTM1 and down-regulation of microtubule-associated protein LC3-II level demonstrating the effects of knockout on phagophore de-novo formation and/or elongation. Accordingly, *atg7* and *beclin1* mutants couldn’t form mature autophagosomes resulted in inhibition of cargo sequestration which accounts for increasing of the p62/SQSTM1 level (39).

Hyperplasia of multilayered columnar epithelium, dysplasia with nuclear atypia and cellular pleomorphic malformations are all phenotypes for aged alimentary of zebrafish (40) which probably be detected in our *atg7* and *beclin1* mutants zebrafish, suggesting that autophagy blocking has a reverse effect on the longevity and accelerates GIT senescence. Furthermore, it has been showed that the embryonic phenotype of *beclin1* null mice is more severe than that *atg5*^−/−^ and *atg7*^−/−^ mice (11) suggesting that *beclin1* regulated early embryogenesis processes more than *atg7*. Consistent with these findings, *beclin1* knock-out zebrafish died earlier than *atg7* null larvae.

Regarding the molecular mechanism, the dynamic cross-talk between autophagy, glycogen and lipid metabolism has been previously studied in *atg7*^−/−^ furrier models that exhibited lipoatrophy with hyperlipidemia and hyperglycemia (41). During starvation, hepatocytes mobilize glycogen stores promptly to increase the availability of glucose and maintain blood glucose and amino acid balance (42). At the same time, it induces lipolysis or in other meaning; it accelerates free fatty acids from peripheral adipose to the liver in order to start the process of ketogenesis then gluconeogenesis (43). Briefly, there is a high flux through fatty acid β-oxidation including the conversion of pyruvate and other anaplerotic substrates into glucose via gluconeogenesis and this has been confirmed in both fasted humans and animal models (44–46).

It has been reported that targeted deletion of essential autophagy genes in mice has various important functions of autophagy including lipid droplet and triglycerides formation (47–49). Together with our observations, indigested lipid droplets accumulated during nutrient deprivation and inhibition of autophagy reflects the essential role of *atg7* and *beclin1* in lipid metabolism (50). Alongside, our results had shown up-regulation of transcription factor *srebf1*, *ACC1*, and *fasn* that are the key enzymes in the lipid de novo formation (51). Besides that, genes involved in lipid β-oxidation including *acox1*, *cpt1aa* were highly declined in mutant strains, as well as, *cd36* has been reported to be increased in non-alcoholic fatty liver disease (52). All data together suggest that lipid accumulation in our autophagy knocked out models is a result of blocked lipolysis.

On the other side, genes involve in glycolysis including *hk1*, *pfk1b*, *gck* were up-regulated in mutants after 5dpf as larvae struggle for surviving in another catabolic pathway (glycolysis) to compensate autophagy impairment. Moreover,(*g6pc3*) enzyme which catalyzes the conversion of glucose 6-phosphate (G6P) to glycogen within the hepatocyte (53) was highly reduced in both mutant strains, unlike wild type that maintains the balance between gluconeogenesis and glycolysis during fasting and feeding transition. All parameters and results indicated the correlation between autophagy and the glycogen boundary equilibrium during the transition from fasting to feeding (14, 54).

Rapamycin might be effective in age-related pathologies via activation of autophagy by enhancing the clearance of aggregate-prone proteins in vitro (55–57) However, Stimulation of autophagy by rapamycin in knocked-out models is still poorly understood. Previously, morphometric analysis of rapamycin-treated zebrafish has been established that revealed a great reduction in the epithelial cells number, size and proliferation hence lower yolk consumption till 9dpf (26, 58). In our study, rapamycin increasing the longevity of *atg7*- and *beclin1*-mutated through slowing metabolism and dietary restriction which reduced cellular food demand without malnutrition resulted in metabolic equilibrium. Nevertheless, rapamycin couldn’t stimulate autophagy in mutants throughout the time of treatment as *atg7* and *beclin1* unable to convert LC3-I intoLC3-II or restore the cargo receptor P62/SQSTM1 which expected to be accountable for the sever deliration of the intestine and strongly correlated with the early death despite treatment (59–61). Those results indicating that *beclin1* and *atg7* are crucial for rapamycin-induced and that rapamycin depends on the *atg5-atg7* dependent autophagy pathway (conventional autophagy) that needs the presence of all ATGs parts including *atg7*as well as *beclin1*.

Briefly, however, the processes that govern success during the transition process from larval to juvenile are complex and not fully understood; autophagy-related genes in our study provided molecular evidence that indicated the role of autophagy during the turning point in the timeline of zebrafish development (Fig.10A). During larval-juvenile transition, perturbations of our studied genes ceased the whole autophagy pathway made the larvae shifted to an easier catabolic pathway and depend on the stored hepatic glycogen, at the same time they unable to utilize lipid and/or start gluconeogenesis. Accordingly, the entire metabolic disturbance affects the structure and function of the gastrointestinal tract and accessory organs and leading to death during larval-to-juvenile transition even though supporting larvae with external food supply (Fig.10B).

**Figure10.**
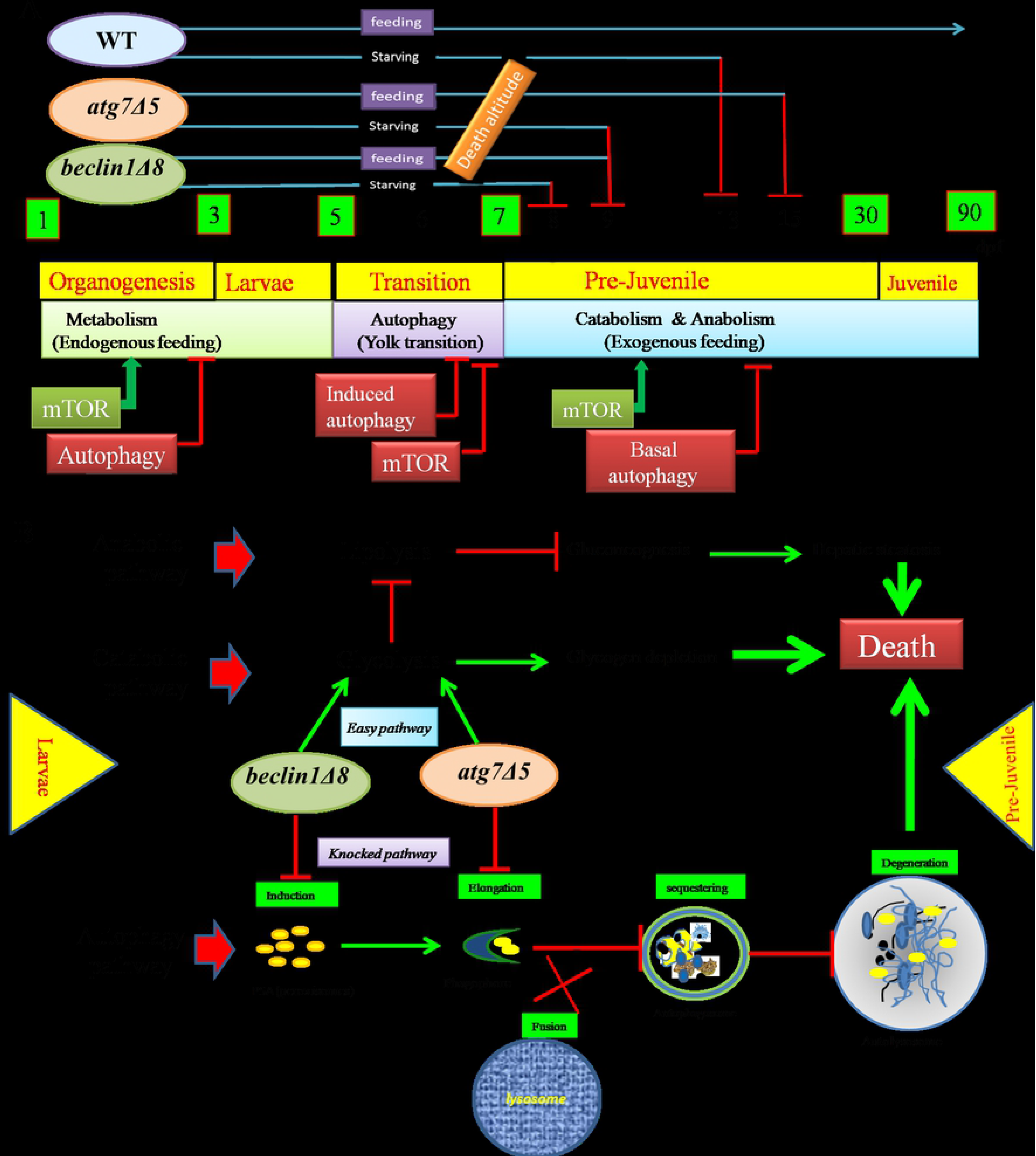
A proposed model of *atg7* and *beclin1* functions during the larvae-juvenile transition. (A): Schematic diagram represents the embryonic development timeline and the observed death altitudes of studied mutants as result of autophagy perturbation. (B): Schematic diagram shows the possible death pathways that affect mutant during early development after yolk depletion indicating the role of autophagy during the shifting from endogenous to exogenous feeding.

## Materials and methods

### Fish husbandry and mutagenesis screen

Zebrafish (*Danio rerio*) AB strain was raised according to the established protocols (62). All experiments involving zebrafish were approved and in compliance with the requirement of the animal care institution and use committee of Huazhong Agricultural University. Adult zebrafish were kept in the recirculating system at 28.5°C with 14 h light/10 h dark cycle and larvae were staged by morphology and age (hours post fertilization, hpf; days post fertilization, dpf).

To generate *atg7*- and *beclin1*-mutated zebrafish, CRISPR/Cas9 vectors were constructed for editing selected specific sites and regions (24). All sgRNAs were designed using CRISPR RGEN Tools (http://www.rgenome.net). The linearized Cas9 plasmids were transcribed into mRNA using the T7 m MESSAGE Kit (Ambition, USA) and gRNA was synthesized using transcript Aid T7 High Yield Transcription Kit (Thermo Scientific, USA). Zebrafish embryos at the one-cell stage were co-injected with 20pg target gRNA and 300pg Cas9 mRNA. For genotyping of the mutant zebrafish, PCR was performed with primers *atg7*-F: 5’-AAATGCCACAGTCCTCCTC-3’, *atg7*-R: 5’-TGAGCCCAGCCTTTATTCT-3’, *beclin1*-F: 5’-GTATGCCATCAACCTCCTA-3’ and *beclin1*-R: 5’-AAAGTGAAGCACTGCGAAT-3’.

### Survival rates, histological assessments, and Whole-mount ORO

For survival rate, after hatching, mutants of both *atg7* and *beclin1* strains separated via in vivo darken liver (100 mutants of both strains required to start the experiment). On the other hand, 100 eggs were collected from wild type mating. We also depended on genotyping of dead embryos every day to make sure of the mutagenesis especially for those treated with rapamycin. The survival probability at any particular time is calculated by the formula given below and curves given by The Kaplan-Meier plot (63).

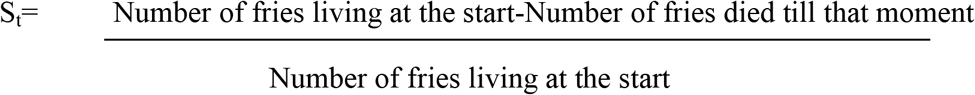

For histological observations, *atg7*-mutated and WT larvae were collected at 5, 7 and 14 dpf, while *beclin1*-mutated were collected before death at 5 and 7 dpf. We opted this time window when the GIT became totally differentiated to follow the digestive architecture during autophagy impairment milestones after yolk transition. Genomic DNA of the tail regions n=48 (+/+: +/−: −/− = 15: 22: 11) was extracted using sodium hydroxide and Tris-HCl buffer at PH 8 (64) and the whole larvae bodies were fixed in 4% paraformaldehyde in phosphate buffered saline (PFA) at 4°C overnight.

Histological sections preparation method achieved as described elsewhere (65). Briefly, the specimens were dehydrated with ethanol series and embedded in paraffin. Tissue sections of 5 μm were prepared using a microtome (Leica Model RM2155). For light microscope analysis, histological sections were stained with hematoxylin and eosin (H&E), periodic acid-Schiff (PAS) and Oil Red O (ORO) as previously described (66).

Whole-mount ORO staining was carried out on 4% PFA fixed larvae. After dehydration in ascending series of 1, 2-propanediol, larvae dyed with 0.5 % Oil Red O solution for 12 h at room temperature then washed in PBS for 20 min and stored at 80 % 1, 2-propanediol till imaging.

### Quantitative real-time PCR (qRT-PCR) and whole-mount RNA in situ hybridization (WISH)

Total RNA was extracted from 35 wild-type larvae using TRIzol (Invitrogen, USA) in successive days from 5dpf to 8dpf. For *atg7* and *beclin1* after genotyping of tail regions, whole body homogenates of 36-38 mutant larvae from both strains were collected stored at −80°C degree till RNA extraction. RNA quality and concentration were evaluated by gel electrophoresis and NanoDrop 2000, respectively. Isolated RNA was reverse transcribed into cDNA was by Prime Script TM RT reagent Kit with gDNA Eraser (Stratagene, Takara). Real-time PCR was performed on ABI-7500 real-time PCR machine (Applied Biosystems, USA) and β-actin was used to normalize the expression values(67).The primer sequences used in qRT-PCR are listed in tables 1, 2, and 3.

For the whole mount in situ hybridization analysis, the partial cDNAs of markers for differentiated hepatocytes liver fatty acid binding protein 10a *fabp10a* (68), intestinal epithelium enterocyte differentiation marker *villin1*(69), pancreatic *insulin* and *trypsin* (70, 71) were amplified using primers listed in table4. Probes were synthesized by in-vitro transcription using DIG labeling mix and T7 polymerase (Roche, USA). Hybridization was conducted as previously described (72). The photograph was taken using Leica MZ16FA Microscope by ACImage software and genotyping was performed after imaging to confirm our results.

### Western blotting and immunohistochemistry assay

In western blot, trunk regions of sequenced larvae tail of *beclin1* n=132 (+/+: +/−: −/− = 32: 63: 37) and *atg7* n=126 (+/+: +/−: −/− = 31: 60: 35) were collected and stored at −80°C. Homogenates from homozygous larvae of each strain were digested using lysis buffer containing a protease inhibitor. Samples were boiled for 10 min at 100°C in a ×5 loading buffer, after that, 15μl of total proteins were subjected to SDS-PAGE (Bio-Rad) and carry out electrophoresis then the fixed gel was electro-transferred it to a nylon membrane. After blocking with 5% skimmed milk in TBST, nylon membranes were incubated with Primary antibodies include rabbit anti SQSTM1/P62 (MBL, PM045), rabbit anti-LC3A/B (Cell Signaling Technology, 4108), and anti-β-actin (Cell signaling, 4967S) respectively. Blots were probed with HRP-conjugated secondary antibody visualized using ECL western blotting detection reagents.NIH software Image J was applied for blot scanning and protein area quantifications.

For immunohistochemistry, fixed embryos at 7dpf were dehydrated in ascending ethanol, embedded in paraffin and sectioned at 5 μm intervals using a Reichert-Jung 2050 microtome (Leica). Sections were deparaffinized and hydrated following by 20 min of antigen retrieval in sodium citrate buffer (pH 6.0) at 100°C. Slides were treated with 0.3% H_2_O_2_ for 10 min to remove endogenous peroxidase then blocked with 5% BSA in PBST for 1 h at room temperature. Sections were incubated with first antibodies, rabbit anti-LC3B (Abcam, ab483940) (1:200) and anti-SQSTM1/P62 (MBL, PM045), (1:500) at 4°C overnight. After 3x PBST washing, 50-100μl HRP secondary antibody was added following staining 3,3′-diaminobenzidine (DAB) substrate and counterstained with hematoxylin for nuclear differentiation.

Immunofluorescence was performed as following steps; after Antigen retrieval by sodium citrate buffer, slides were blocked with PBS with TritonX-100(PBT) containing 5 % BSA and for 1 hour, then incubated with primary antibodies mentioned previously diluted in PBT overnight 40°C. Washing three times again in PBST and incubated with fluoresce in-conjugated secondary antibodies for 1 hour at room temperature. Nuclei were stained with 4′, 6-diamidino-2-phenylindole (DAPI) and mounted. Sections were analyzed by fluorescence microscopy using an Axio Vision image capture system.

### Drug treatment

WT, *atg7* and *beclin1*-mutated embryos were treated from 1st till 7^th^ dpf in embryo-medium at 28.5 °C with 400nM rapamycin (CAS53123-88-9 MedChem, Express) prepared in dimethyl sulfoxide (DMSO). 0.1% DMSO solution (the treatment vehicle) was used for the control treatment. Drug-containing media were replaced every 24 h. Larvae were collected at 7dpf where tails genotyped and trunk regions were fixed in a mixture of 40% ethanol, 5% acetic acid and 10% formalin all night at 4°C for immunohistochemistry, and 4% PFA for histological observations. Nearly 35-40 genotyped larvae from control and each treated group were used for protein extraction and blotting assay in the western blot experiment.

### Statistical analysis

Data are presented as mean ± SD (n=3). Statistical analyses were performed using SPSS and the data were analyzed by student t-test and one-way ANOVA (*P < 0.05;**P < 0.01; ***P < 0.001). Plots were designed using graphpad prism 8 software.

## Acknowledgments

This work was supported by the Fundamental Research Funds for the Central Universities (534-180010235, 2662017PY013). The funders had no role in study design, data collection, and analysis, decision to publish, or preparation of the manuscript.

## Supporting information legends

### 1 Supporting figures legends

**S1 Fig: HSP distribution on query sequence of *atg7* and *beclin1* CRISPR/Cas9 knock-out target sites.** (A): Sequences information of *atg7* target site blast against zebrafish genome. (B): The location information of target sites *atg7* in chromosome 11 in zebrafish genome. (C): Sequences information of *beclin1* target site blast against zebrafish genome. (D): The location information of *beclin1* target sites in chromosome 12 of zebrafish genome. The red box represents the location of the target gene while the red arrow represents the location of target site. According to the information of zebrafish *atg7* and *beclin1* in Ensembl (atg7 ENSDARG00000102893, becn1 ENSDARG00000079128).

**S2 Fig: Agarose gel electrophoresis of Cas9 mRNA and gRNA mRNA.** (A): Agarose gel electrophoresis of Cas9mRNA, marker used is DL2000 DNA marker, L1, L2, L3 indicates Cas9 mRNA in triplicate running. (B): Agarose gel electrophoresis of *atg7* gRNA using DL2000 DNA marker, lines L1,L2,L3 indicates gRNA in triplicate running with 150bp. (C): Agarose gel electrophoresis of *beclin1* gRNA using DL2000 DNA marker, lines L1,L2,L3 indicates gRNA in triplicate running with 200bp.

**S3 Fig: Schematic diagram depicting the functionality of the CRISPR/Cas9 system**. (1): Humanized Cas9 and gmRNA were co-injected in one cell wild type eggs.(2): Confirmation of target mutation by PCR and PMD cloning from the tail region of the grown injected eggs.(3): Generation of F1 from paired mating of heterozygous and wild type strains. (4): Generation of F2 from heterozygous of the same population mating to produce 25%. of mutant embryos with the effective gene knockout.

**S4 Fig: Agarose gel electrophoresis of PCR amplification results**. Genomic DNA was extracted from the larvae tail and PCR was conducted at 58°c annealing temperature. The product length was 399bp in *atg7* and 310 for *beclin1* and most of our results were effective appeared as bright band at right amplification using DL 2000 DNA marker.

**S5Fig: Genotyping results of the PCR amplifications of all predicted larvae WT, Heterozygous and mutant strains within the same population.** (A): Result of *atg7* gene mutation detection. The red box represents the target sit of *atg7* gene and the blue arrow represents the actual mutations in heterozygous by reverse primer. (B): Result of *beclin1* gene mutation detection. The red box represents the target site and the blue arrow represents the actual mutations in heterozygous by the forward primer

### 2 Supporting tables legends

S1Table: Primers used in qRT-PCR (Expression of autophagy-related genes).

S2 Table: Primers used in qRT-PCR (Expression of genes involved in glycogen metabolism).

S3 Table: Primers used in qRT-PCR (Expression of genes involved in lipid metabolism).

S4Table: Primers used in cDNA amplification for probe synthesize (WISH).

S5Table: Primer sequences of *atg7* and *beclin1* gRNA template used in our zebrafish model.

